# Single-cell analysis of Rohon-Beard neurons implicates Fgf signaling in axon maintenance and cell survival

**DOI:** 10.1101/2023.08.26.554953

**Authors:** Adam M. Tuttle, Lauren N. Miller, Lindsey J. Royer, Hua Wen, Jimmy J. Kelly, Nicholas L. Calistri, Laura M. Heiser, Alex V. Nechiporuk

## Abstract

Peripheral sensory neurons are a critical part of the nervous system that transmit a multitude of sensory stimuli to the central nervous system. During larval and juvenile stages in zebrafish, this function is mediated by Rohon-Beard somatosensory neurons (RBs). RBs are optically accessible and amenable to experimental manipulation, making them a powerful system for mechanistic investigation of sensory neurons. Previous studies provided evidence that RBs fall into multiple subclasses; however, the number and molecular make up of these potential RB subtypes have not been well defined. Using a single-cell RNA sequencing (scRNA-seq) approach, we demonstrate that larval RBs in zebrafish fall into three, largely non-overlapping classes of neurons. We also show that RBs are molecularly distinct from trigeminal neurons in zebrafish. Cross-species transcriptional analysis indicates that one RB subclass is similar to a mammalian group of A-fiber sensory neurons. Another RB subclass is predicted to sense multiple modalities, including mechanical stimulation and chemical irritants. We leveraged our scRNA-seq data to determine that the fibroblast growth factor (Fgf) pathway is active in RBs. Pharmacological and genetic inhibition of this pathway led to defects in axon maintenance and RB cell death. Moreover, this can be phenocopied by treatment with dovitinib, an FDA-approved Fgf inhibitor with a common side effect of peripheral neuropathy. Importantly, dovitinib-mediated axon loss can be suppressed by loss of Sarm1, a positive regulator of neuronal cell death and axonal injury. This offers a molecular target for future clinical intervention to fight neurotoxic effects of this drug.

## MATERIALS & METHODS

### Zebrafish husbandry

Adult zebrafish were maintained at 28.5°C as previously described (Westerfield 2000). Embryos were derived from natural matings or *in vitro* fertilization, raised in embryo media, and developmentally staged (Kimmel et al. 1995). Strains utilized were *AB, *SAIGFF213A;UAS:GFP* (“*RB:GFP”) (Muto et al. 2011)*, *sarm1^hzm13^* (Tian et al. 2020), *hsp70:dnFGFR1a-GFP* (Lee et al. 2005), *isl1[ss]:LEXAV-P16;LEXAOP:tdTomato (Rasmussen et al. 2015), dusp6:gfp (Molina et al. 2007)*, and TgBAC(*neurod1:EGFP*)*^nl1^*(Obholzer et al. 2008). The developmental stages utilized in this work are prior to sex specification and animals used in experiments are of indeterminate sex.

### Genotyping protocols

The *sarm1^hzm13^* allele (Tian et al. 2020) was detected by PCR based genotyping using the following primers: Forward: 5′-GATTTGCCGTTATCTCTCCA-3′; Reverse: 5′-TCAAGCAGTTTGGCAGACTC-3′. PCR products were digested with Aat II (New England Biolabs) for 4 h at 37° C, and run out on a 2% agarose TBE gel to resolve digested bands. Wild-type allele is digested into 277 and 129 bp fragments, while mutant allele remains uncut.

### Larval heat-shock protocol

Larvae were transferred to 8-strip PCR tubes in 100 μL embryo medium (2-3 larvae/tube), and then ramped to 37° C for 30 min in a thermocycler, then immediately transferred to 28.5° C embryo medium to recover.

### Embryo dissociation and FACS

Single cell suspensions from day 7 spinal cords were prepared as previously described. For 30 hpf data set, TgBAC(*neurod1:EGFP*)*^nl1^*zebrafish embryos were collected and euthanized in 1.7 ml microcentrifuge tubes. Embryos were deyolked using a calcium-free Ringer’s solution (116 mM NaCl, 2.6 mM KCl, 5 mM HEPES pH 7.0), by gently pipetting up and down with a P200 pipet. Embryos were incubated for 5 minutes in Ringer’s solution. Embryos were transferred to pre-warmed protease solutions (0.25% trypsin, 1 mM EDTA, pH 8.0, PBS) and collagenase P/HBSS (100 mg/mL) was added. Embryos were incubated at 28° C for 15 minutes and were homogenized every 5 minutes using a P1000 pipet. Stop solution (6X, 30% calf serum, 6 mM CaCl_2_, PBS) was added and samples were centrifuged (350xg, 4° C for 5 minutes). Supernatant was removed and 1 mL of chilled suspension solution was added (1% FBS, 0.8 mM CaCl_2_, 50 U/mL penicillin, 0.05 mg/mL streptomycin, DMEM). Samples were centrifuged again (350g, 4° C for 5 minutes) and supernatant was removed. 700 μl of chilled suspension solution was added and cells were resuspended by gentle pipetting. Cells were passed through a 40 μm cell strainer into a FACS tube and kept on ice. GFP+ cells from TgBAC(*neurod1:EGFP*)*^nl1^* zebrafish embryos were sorted for the brightest 2% of GFP+ cells to enrich for sensory neurons on a BD Symphony cell sorter into sorting buffer (50 μl PBS/ 2% BSA) in a siliconized 1.5 mL tube.

### 10X chromium scRNA-seq library construction (30 hpf neurod:egfp data set)

Approximately 15,000 cells were loaded into the Chromium Single Cell Controller to generate barcoded RT products (Chemistry Ver 3.1; 10X Genomics, Pleasanton, CA, USA). The library was sequenced using Illumina NovaSeq 6000 to a depth of at least 40,000 reads per cell.

### Quality Control and Unsupervised clustering

Data sets, single cell reads were aligned to a modified Ensembl version GRCz11 of the zebrafish genome (Lawson et al. 2020) by the Integrated Genomics Laboratory (Oregon Health & Science University) using Cell Ranger (version 6.1.1; 10X Genomics, Pleasanton, CA. USA). The UMI count matrix was analyzed using Seurat (version 4.0.1) (Butler et al., 2018). Quality control filtered out genes expressed in fewer than three cells, cells with less than 1,900 unique genes, and cells that expressed greater than 5% of mitochondrial transcripts. The remaining cells subjected to further analysis. Linear dimensionality reduction, clustering and UMAP visualization were performed with Seurat (Butler et al., 2018). For the spinal cord data set, principal component analysis was performed to project the 2,000 most variable genes into 30 principal components. Thirty clusters were identified with the Seurat implementation of the Louvain algorithm using a resolution of 1, and visualized with 2 UMAP dimensions. RB clusters were identified using the following markers: *isl2a*, *prdm14*, and *fgf13a*.

For the data set derived from 30 hpf *TgBAC(neurod:egfp)^nl1^*embryos, 5,235 cells were subjected to analysis after QC. Principal component analysis was performed to project the 2,000 most variable genes into 30 principal components. A total of 36 clusters were identified with the Seurat implementation of the Louvain algorithm using a resolution of 2 and visualized with 2 UMAP dimensions. Clusters were manually annotated using a whole zebrafish single-cell transcriptome atlas (Farnsworth et al. 2020) as well zebrafish database of gene expression (ZFIN expression atlas: (Thisse 2001)). Two clusters containing trigeminal and Rohon-Beard neurons were identified using known markers, including *isl2b*, *kitb*, *prdm14*, *trpv1*, *prdm14*, and *fgf13a*. These were subset and reclustered using 20 principal components and a resolution of 1. Three new clusters were identified that roughly correspond to the same three RB clusters identified using neurons from the trunk (*kitb*+, *tmem178*+, and *adcyap1a*+).

Classification of head- and trunk-specific cells was performed using expression of known posterior *hox* genes. The gene *hoxd3a* was chosen for assignment of the posterior identity because it is expressed throughout the hindbrain and spinal cord caudal to rhombomere 5. Seurat’s WhichCells function was used to classify “*hoxd3a*+” and “*hoxd3a*-” cells with a scaled expression level cutoff of 0.35; Seurat’s FindMarkers function was then used to identify differentially expressed genes between these two subsets. To compare zebrafish RB with mouse reference data set we used biomaRt (version 2.54.0) and the reference genomes GRCz11 (zebrafish) and GRCm39 (mouse). We than used 50 top DE genes for each RB subtype to generate Module Scores using AddModuleScore Seurat function. To generate Pearson correlation coefficients between zebrafish RB subtypes and subtypes of mouse TG and DRG neurons, we identified 2000 Variable features of all zebrafish RBs. These filtered down to the ∼1300 mapped mouse homologs. Expression levels of these genes in each subcluster were used to calculate correlation coefficients.

### Drug treatment

Dovitinib and SU5402 were obtained (Selleck Chemicals, Houston, TX, USA) and dissolved in DMSO to stock solutions of 10 mM and stored at -80° C. For larval treatment, drugs were diluted and mixed in embryo medium containing 1% DMSO and used to replace larval media. Unless otherwise noted, larvae were treated with drugs from 72-120 hpf and larval media was replaced at 96 hpf with fresh drug/vehicle-containing media.

### Zebrafish fluorescent in situ hybridization and immunostaining

For FISH, we employed commercial probes designed by Advanced Cell Diagnostics (ACD) according to a published protocol (Gross-Thebing 2020) with following modifications. We used RNAscope Multiplex Fluorescent Reagent Kit (ACD). Zebrafish larvae were fixed overnight at 4°C in BT-fix (4% PFA in PBS with 4% sucrose and 0.15 mM CaCl_2_). After fixation and wash in PBST (1xPBS+0.01% Tween-20), larvae were dehydrated through a methanol wash series (25, 50, 75% MeOH in PBST for 5 minutes each) then stored at - 20°C overnight. Before protease digestion, larvae were rehydrated through the same methanol wash series (75, 50, 25% MeOH in PBST for 5 minutes each) and rinsed with PBST for 5 min. v2RNAscope kit utilizes an HRP/tyramide dye-based detection system as follows. HRP-C1 was added to larvae and they were incubated at 40°C for 15 minutes, then washed in SSC-Tween 0.01% 3×15 minutes at room temperature with gentle agitation. Opal 570 (Akoya Biosciences) was diluted 1:1000 in TSA buffer (Advanced Cell Diagnostics), added to larvae, and incubated at 40°C for 30 minutes, then washed as above. Larvae were incubated in HRP blocker for 15 minutes at 40°C then washed as above. These steps were repeated with HRP-C2 and Opal 690 (Akoya Biosciences), then with HRP-C3 and CF405M (Biotium). For the detection of GFP signal, embryos were post-fixed with 4% paraformaldehyde in 1× PBS for 20 min at room temperature (RT), then processed according to the protocol described below.

For immunolabeling, larvae were anesthetized in 0.02% tricaine (MS-222; Sigma) in embryo medium, fixed in 4% paraformaldehyde in 1× PBS for 1 hour at RT and then 4°C overnight. Larvae were rinsed 3x with 1× PBS and permeabilized by washing in distilled water for 5 min, followed by 3x washes in 1× PBS + 0.1% Triton X-100 (PBTx). Larvae were incubated for 1 hour at RT in blocking solution (PBTx + 5% goat serum, 1% bovine serum albumin, 1% DMSO), then at RT in primary antibody diluted in blocking solution for 2 h followed by 4° C overnight. Embryos were rinsed extensively in PBTx and incubated for 4 hours at RT in Alexa488-, Alexa568-, or Alexa647-conjugated secondary antibodies diluted in blocking solution (1:1000, ThermoFisher Scientific). After rinsing in PBTx, larvae were transferred to 70% glycerol in PBS and mounted on slides with 0.13-0.17 mm borosilicate coverslips secured with vacuum grease. Primary antibodies used were α-GFP (1:1500, Aves Labs, RRID:AB_10000240) and α-Cleaved Caspase-3 (1:1000, Cell Signaling, 9664P, RRID:AB_2070042).

### Live imaging

For live-imaging, larvae were mounted in 1.5% low melting point agarose on a glass coverslip, submerged in embryo media containing 0.02% tricaine (MS-222; Sigma), and imaged using a 40x/NA=1.25 silicon oil immersion objective on an upright Fluoview3000 confocal microscope (Olympus) with 488 and 568 nm excitation channels. For dovitinib treatment time-lapse, we cut away agarose around the head and tail to allow better drug access. Z-stacks were collected approximately every 5-8 min for 10-12 hours. For time-lapse imaging of RB neuron cell bodies, the anterior-most edge of the 360 μm viewing window was placed in line with the end of the yolk extension.

### Confocal imaging of FISH and immunolabeled samples

Stained larvae were imaged on the Olympus Fluoview3000 with 405, 488, 568, and 647 nm excitation channels using a 20x/NA=0.75 air objective. We imaged two 400 μm windows immediately proximal and distal to the end of yolk extension. As we did not observe any significant differences in marker expression or Caspase-3 labeling between these two regions, we combined counts for each individual. RB neurons were identified by spinal cord location, morphology, and axonal projections. Counts per viewing window were normalized to a 100 μm length.

### High-throughput fluorescence larval imaging

For live high-throughput imaging of caudal tails of wild-type and dovitinib-treated larvae we used the Large Particle (LP) Sampler and VAST BioImager (Union Biometrica) to automate the delivery of zebrafish from 96 multi-well plates to the BioImager microscope platform as previously described (Tuttle et al. 2022). Z-stacks of 218 frames with a step size of 1.84 μm were acquired using Zen Blue 2.0 software (Carl Zeiss).

### Axon density quantification

Caudal tail axon density was measured as previously described (Tuttle et al. 2022). Briefly, we imported confocal images of *RB:GFP* caudal tails into FIJI (Schindelin et al. 2012), converted images to 8-bit, and generated maximum intensity Z-projections. We then thresholded the projections, formed a rectangular 50×100 μm ROI at the distal tail tip, and measured the percentage of this ROI occupied by GFP+ axons.

### Experimental design and statistical analysis

The sex of the animals was unknown as sex specification has not occurred at this stage of larval development. Analysis was performed with Prism software (Graphpad). Specific statistical tests and post hoc tests for each data set are indicated in text and figure legends. For experiments involving more than two independent variables, ANOVA tests were performed to test if main effects and interactions were statistically significant. If interaction was statistically significant, ANOVA was repeated with simple effects, and significance of main effects was re-evaluated and the indicated post hoc tests were performed. Results were considered statistically significant when *p* < 0.05. Data in text or plotted with error bars on graphs were expressed as mean ± SEM.

### Image Processing

Images were processed using ImageJ (Abramoff et al., 2004; Schindelin et al., 2012) or Imaris (Bitplane) software.

### Data Access

scRNA-seq data are publicly available through Gene Expression Omnibus (GEO) database: 4 dpf spinal cord data sets accession number is GSE232801; 7 dpf spinal cord data set accession number is GSE241296; and 30 hpf TgBAC(*neurod1:EGFP*)*^nl1^*data set accession number is GSE240721. The code used for scRNA-seq data analyses as well as to generate Venn diagrams is available on Github: https://github.com/anechipor/nechiporuk-lab-Tuttle_et_al_2023.

### Study Approval

All animal works were approved by and conducted according to guidelines of the Oregon Health and Science University Institutional Animal Care and Use Committee, protocol# IP00000495.

## INTRODUCTION

Rohon-Beard neurons (RBs) are a population of somatosensory neurons present in amniotes during embryonic, larval, and juvenile stages. RBs are located in the dorsal spinal cord and respond to mechanical, thermal, or chemical stimuli (Metcalfe et al. 1990; Prober et al. 2008; Low et al. 2010; Gau et al. 2013; Ogino et al. 2015). RBs have become a popular model for screening drug neurotoxicity as well understanding axon damage and axon regeneration (Rieger and Sagasti 2011; Cuevas et al. 2013; Doganli et al. 2013; Wang et al. 2015; Adula et al. 2022). In particular, zebrafish RBs have been used to study the cellular and molecular bases of drug induced peripheral neuropathies (DIPNs) (Lisse et al. 2016; Cadiz Diaz et al. 2022; Tuttle et al. 2022). DIPNs are caused by damage to the peripheral sensory nervous system as an unintended, off-target effect from many different types of small molecular drugs to treat cancers, including both cytotoxic and targeted therapeutics (*e.g*. multi-kinase inhibitors or MKIs) (Peltier and Russell 2002; Visovsky 2003; Kane et al. 2006; Breccia and Alimena 2010; Cortes et al. 2010; Rey et al. 2015; Roy et al. 2019; Merheb et al. 2022). Treatment options for patients experiencing painful DIPNs are highly limited and non-specific (Shah et al. 2018). Thus, there is an urgent need to identify particular targets and downstream effectors that cause this neurotoxicity. RBs offer a powerful model system to identify and explore these pathways, but it is important to understand whether RBs represent a molecularly heterogenous population of neurons, similar to dorsal root ganglia (DRG) and trigeminal (TG) ganglion neurons.

RBs are induced during gastrulation at the neural plate border and become functional by 48 hours post-fertilization (hpf) (Rossi et al. 2009). RB neurons remain functional for weeks until juvenile stages when they become replaced by DRGs as the primary somatosensory neurons (Rasmussen et al. 2018). Previous studies describe multiple subtypes of RB neurons (Slatter et al. 2005; Pineda et al. 2006; Appelbaum et al. 2007; Patten et al. 2007; Pan et al. 2012). Palanca and colleagues used multiple transgenic reporters with enhancers to visualize gene expression in RBs (Palanca et al. 2013) and concluded RBs fall into at least three overlapping categories: *p2rx3b*-positive; *pkca*-, *p2rx3a*-, and *trpA1b*-positive; and *ntrka*-positive. One limitation of their approach is that it utilized a small number of existing and new reporter lines. Thus, a more unbiased, transcriptionally-based approach is warranted.

In this study, we utilized scRNA-seq to define three distinct, non-overlapping populations of RBs present in zebrafish at larval stages. Single-molecule fluorescent in situ hybridization (FISH) confirmed the presence of these non-overlapping subtypes. We then used expression of posterior *hox* genes to distinguish populations of TG and RB neurons in our scRNA-seq data set. Our analysis revealed a small number of differentially expressed genes between TG and RB populations, one of which was validated by FISH. To demonstrate the utility of our data set, we tested the requirement for a particular receptor-tyrosine kinase (RTK) pathway – Fibroblast Growth Factor (Fgf) – in RB neurons. We show that several direct Fgf transcriptional targets, as well as an Fgf reporter, are expressed in RB neurons. Furthermore, pharmacological or genetic inhibition of the Fgf pathway leads to a dramatic loss of peripheral RB axons and RB cell death. We also found that application of an FDA-approved RTK inhibitor, dovitinib, which is known to target the Fgf pathway and cause DIPNs, induces identical somatosensory axon loss. Importantly, this phenotype can be suppressed by the loss of *sarm1*, a positive regulator of neuronal cell death and axonal injury, offering a molecular target for future clinical intervention to fight neurotoxic effects of this drug.

## RESULTS

### Identification and transcriptional profile of somatosensory neurons

Previous studies provided evidence for distinct subpopulations of RB neurons based on differences in both electrophysiology and expression of transgenic reporters (Won et al. 2011; Palanca et al. 2013). However, the number and specific identity of these subpopulations have not been definitively explored. Advances in single-cell RNA-sequencing (scRNA-seq) now allow an unbiased approach to classify neuronal subpopulations based on their transcriptional profile. However, RBs make up a small fraction of neurons in a zebrafish larva and most data sets do not have sufficient numbers of RBs sampled to investigate potentially subtle transcriptional differences (Lencer et al. 2021). In addition, RB neurons are closely related to trigeminal neurons, which further complicates their characterization on a transcriptional level. Recent scRNA-seq data sets from isolated neurons in the larval zebrafish spinal cord identified an RB-specific cluster (Kelly et al. 2023). This new resource offers a unique opportunity to investigate possible RB subpopulations compared to less-focused data sets because they: 1) use spinal cords instead of whole larvae further enriching the proportion of RB cells sequenced; and 2) exclude the more anteriorly located trigeminal population that can confound analysis of RBs.

To complement the published day 4 spinal cord data sets, we also isolated and sequenced cells from day 7 spinal cords using the same method (Kelly et al. 2023). These three data sets were subjected to a quality control and unbiased clustering using the Seurat pipeline (6,949 cells) (Fig. 1A). Using published day 4 clustering analysis, we identified two large subpopulations, containing neurons and glia (Fig. 1B,C). RB neurons were identified as clusters that expressed known RB markers such as *isl2a* and *prdm14* (Clusters 24 and 29; Fig. 1D,E). Presumptive RB subclusters were then subjected to another round of unsupervised clustering, which yielded three distinct subpopulations (Fig 1. F-I). To identify genes that are enriched in each population, we performed a differential expression (DE) analysis with a minimum difference expression threshold set at 25% and FDR at <5%. This analysis yielded between 102 and 177 genes specific for each subcluster. Figures 1J-M illustrate top differentially expressed genes (genes that are subsequently mentioned in the text are underlines). Examples of cluster-specific genes included: *kitb* and *runx3* for Subcluster 0 (termed *kitb*+ cells); *calca* and *tmem178* for Subcluster 1 (termed *calca*+ cells); *adcyap1* and *prickle1a* for Subcluster 2 (termed *adcyap1*+ cells).

**Figure 1:**
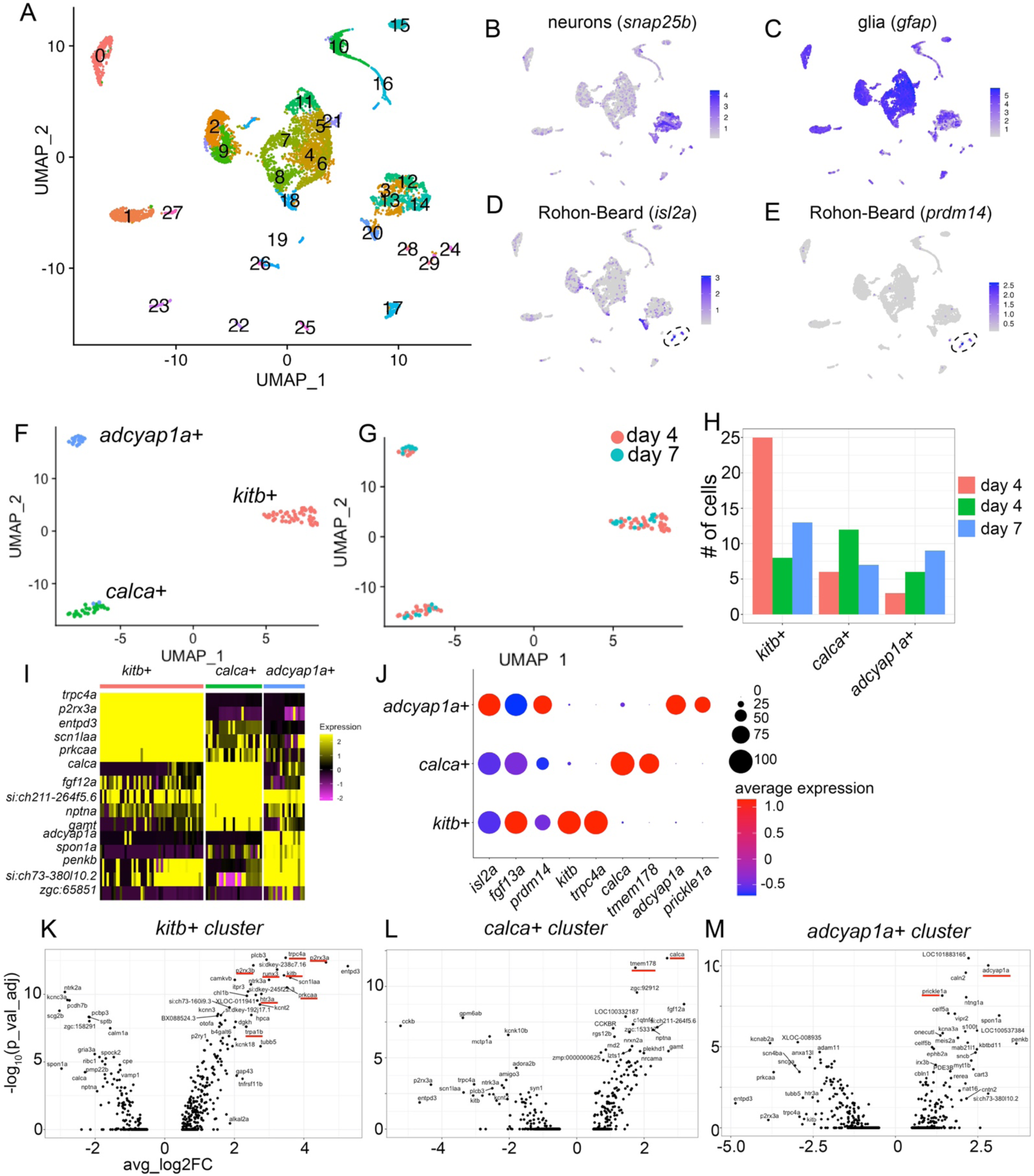
Identification of transcriptionally-distinct RB subpopulations. (A) Unsupervised clustering and UMAP reduction diagram of cells derived from trunks of day 4 (Kelly et al. 2023) and day7 larvae. (B,C) Feature plots identifying neurons (B) and glia (C). (D,E) Feature plots of two RB markers, *isl2a* and *prdm14*, identifying clusters 24 and 29 as RBs. (F,G) Unsupervised subclustering and UMAP reduction diagram of RB cells. (H) Bar plot showing cell numbers for each cluster. (I) Heat map showing top five DE genes for each cluster. (J) Dot plot profile of genes expressed in all RBs (*isl2a*, *fgf13a*, *and prdm14*), *kitb*+ cluster (*kitb* and *trpc4a*), *calca*+ cluster (*calca* and *tmem178*), and *adcyap1a*+ cluster (*adcyap1a* and *prickle1a*). (L-M) Volcano plots showing top DE genes in each RB cluster.

### Three distinct subpopulations of RB neurons

To validate subcluster-specific markers, we performed whole-mount, single-molecule FISH on 3-day old larvae, using combinations of 3 probes: *kitb*+*calca*+*adcyap1a* and *runx3*+*prickle1a*+*tmem178* (Fig. 2). To visualize RBs, we took advantage of a transgenic zebrafish that marks most RBs: a *SAIGFF213A* GAL4 driver (Muto et al. 2011) and *UAS:GFP*; this double transgenic hereafter referred to as “*RB:GFP*”. In this enhancer trap line, the driver is integrated into the *prdm14* locus (Muto et al. 2011), which is expressed in the majority of RBs (Fig. 1E). Consistent with our scRNA-seq analysis, we observed highly exclusive expression of markers of each subcluster. Within the same animal, RBs express *kitb, calca*, or *adcyap1a* and exhibit very little to no overlap (Fig. 2A,B). Similarly, probes for *runx3*, *prickle1a*, and *tmem178* showed minimal overlap (Fig. 2C,D). This *in situ* analysis confirms our scRNA-seq-based identification of three distinct subpopulations of RB neurons.

**Figure 2:**
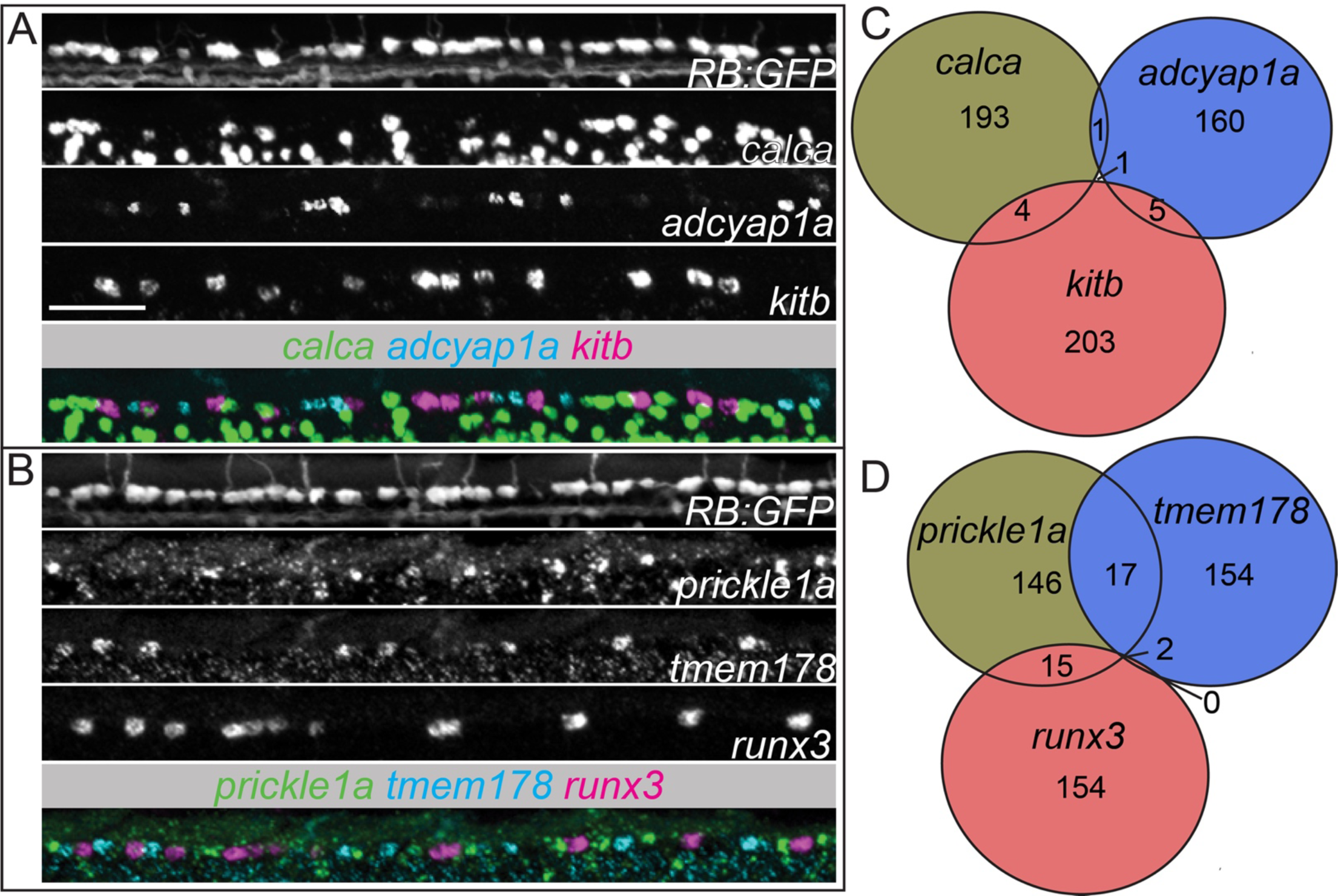
Marker expression for individual RB neuron subpopulations is largely exclusive. (A,B) Lateral view of triple fluorescence *in situ* hybridization (FISH) in *RB:GFP*+ RB neurons in a single 72 hpf larva. (A) RNAscope for *kitb*, *adcyap1a*, and *calca* and (B) *prickle1a*, *tmem178*, and *runx3* mark largely non-overlapping sets of RBs. (C,D) Summed counts of RB markers from the triple FISH experiments. Scale bar=50 μm.

### Differentiating trigeminal from RB neurons

TG and RB neurons are both somatosensory neurons that innervate the larval skin. They arise at the same time, express the same transgenic reporters, and their peripheral neurites have similar branching patterns and growth cone morphology (Metcalfe et al. 1990; Palanca et al. 2013). However, these populations also have notable differences. RB neurons are present in the trunk and arise from the neural plate border (NPB) (Cornell and Eisen 2000). In contrast, TG neurons are located in the head and are formed from both placodal and neural crest cells (Hamburger 1961; Lwigale 2001). However, it is unclear how similar TG and RB subtypes are and whether these populations can be distinguished on a molecular level.

In order to identify markers that are present in only one of the two neuronal populations, we set out to obtain an additional scRNA-sequencing data set containing both TG and RB neurons. To achieve this, we isolated zebrafish neurons using FACS from TgBAC(*neurod1:egfp*)^nl1^ embryos at 30 hpf. Initial subclustering yielded three populations that contained both TG and RB neurons based on common markers; however, it did not distinguish them (Fig. 3A-F). In order to separate these two populations, we took advantage of multiple *hox* genes known to be expressed in the posterior region of the animal (*i.e*. present in RB, but absent in TG neurons), including *hoxc3a*, *hoxd3a*, *hoxa9b*, and *hoxb9a*. Using this approach, we identified several candidate genes that were differentially expressed between RB and TG neurons (Fig. 3H-J). We then examined expression of one of these genes, *heparan sulfate (glucosamine) 3-O-sulfotransferase 1-like1* (*hs3st1l1*), by FISH in RB*:GFP* larvae. Compared to *kitb*, which is expressed in subpopulations of both RBs and TGs, *hs3stl1l* is only present in a subset of RBs, but completely absent from TGs within the same embryo (Fig. 3K, L). This provides an important transcriptional marker that can be used in future analyses of differences between RBs and TGs.

**Figure 3:**
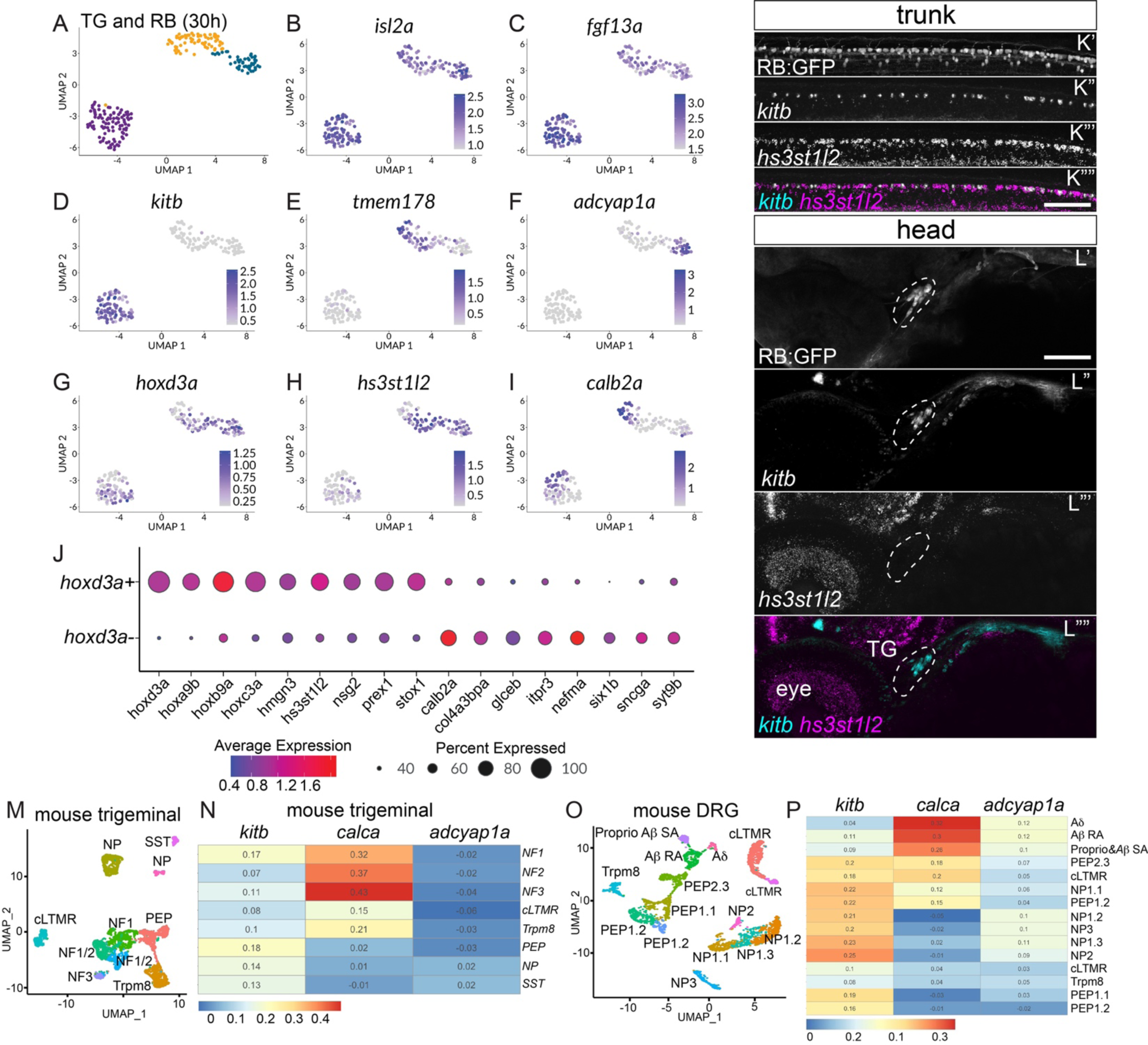
Comparison of RB and TG transcriptional profiles. (A) Unsupervised clustering and UMAP reduction diagram of TG and RB cells derived from 30 hpf TgBAC(*neurod1:egfp)^nl1^*embryos yield three cell clusters containing both TG and RB cells. (B,C) Feature plots demonstrating *isl2a* and *fgf13a* mark both TG and RB cells. (D-F) Feature plots showing presence of RB markers in RB and TG cells. (G) Expression of posterior *hox* gene, *hoxd3a*, marks RB cells. (H) Feature plot of an RB-specific marker, *hs3st1l2*. (I) Feature plot of a TG-specific marker *calb2a*. (J) Dot plot showing expression of posterior hox genes as well as RB-specific markers (*hoxd3a*+ subset) and TG-specific markers (*hoxd3a*-subset). (K,L) Lateral views of FISH staining for *kitb* and *hs3st1l2* at 72 hpf in RB (K) or TG (L) neurons marked in *RB:GFP* larva. Note that subpopulations of RB neurons are positive for *kitb* or *hs3st1l2* while no TG neurons express *hs3st1l2*. Dashed lines=TG. (M) Unsupervised clustering and UMAP reduction diagram of adult mouse TG cells (Yang et al. 2022). (N) Heatmap illustrating module scores for each RB cluster that are associated with each type of mouse TG neurons. Note the *calca*+ RB cluster is most similar to NF3 TG neurons. (O) Unsupervised clustering and UMAP reduction diagram of adult mouse DRG cells (Jung et al. 2023). (P) Heatmap illustrating module scores for each RB cluster that are associated with each type of mouse DRG neurons. Note the *calca*+ RB cluster is most similar to A8 DRG neurons. Scale bar=100 μm

The presence of multiple RB-specific markers implied that these neurons are transcriptionally distinct from other types of peripheral sensory neurons, such as TG and DRG neurons. To further investigate this question, we compared each subtype of RB neurons from day 4 and 7 data sets to published transcriptional profiles of adult mouse TG and DRG neurons (Yang et al. 2022; Jung et al. 2023). We used the top 50 differentially upregulated genes for each RB subtype to generate a Module Score for each of the TG and DRG neuron classes (Seurat function AddModuleScore). There are 8 and 17 transcriptionally distinct classes of TG and DRG neurons, respectively. According to this analysis, the *calca*+ subtype was mostly similar to A-fiber LTMR types of TG (NF3 subtype; Module Score = 0.43; range: -0.36 to 0.92) as well DRG (Aδ subtype; Module Score = 0.32, range: -0.35 to 0.80) neurons (Fig. 4M-P). Interestingly, NF3 TG neurons are highly transcriptionally similar to Aδ DRG neurons (Yang et al. 2022), which is consistent with the above observation. The *kitb*+ subtype of RBs was weakly homologous to peptidergic (PEP) and non-peptidergic (NP) neurons (TG PEP neurons Module Score = 0.18, range: -0.21 to 0.49; DRG NP2 neurons Module Score = 0.25; range: -0.23 to 0.52), whereas *adcyap1a*+ RB neurons did not display any transcriptional similarity to TG or DRG neurons (Fig. 4M-P). As an alternative method, we calculating Pearson correlation coefficients between top ∼1,300 variable genes in RBs that are homologous to mouse for each RB cluster versus TG and DRG neurons. This approach revealed a moderate correlation between the neuronal subtypes that displayed the highest Module Scores: NF3 TG vs. *calca*+ RBs is 0.34, whereas correlation between Aδ DRG and *calca*+ Rb is 0.29. Similarly the correlation between TG PEP vs. *kitb*+ RBs is 0.31 and DRG NP2 vs. *kitb*+ RBs 0.26. In summary, these observations argue that three subclasses of RB neurons are for the most part distinct from the known subtypes of mammalian DRG and TG neurons; although, *calca*+ neurons share some transcriptional similarity with cutaneous mechanoreceptors and nociceptors of group A fibers in TG and DRG.

**Figure 4:**
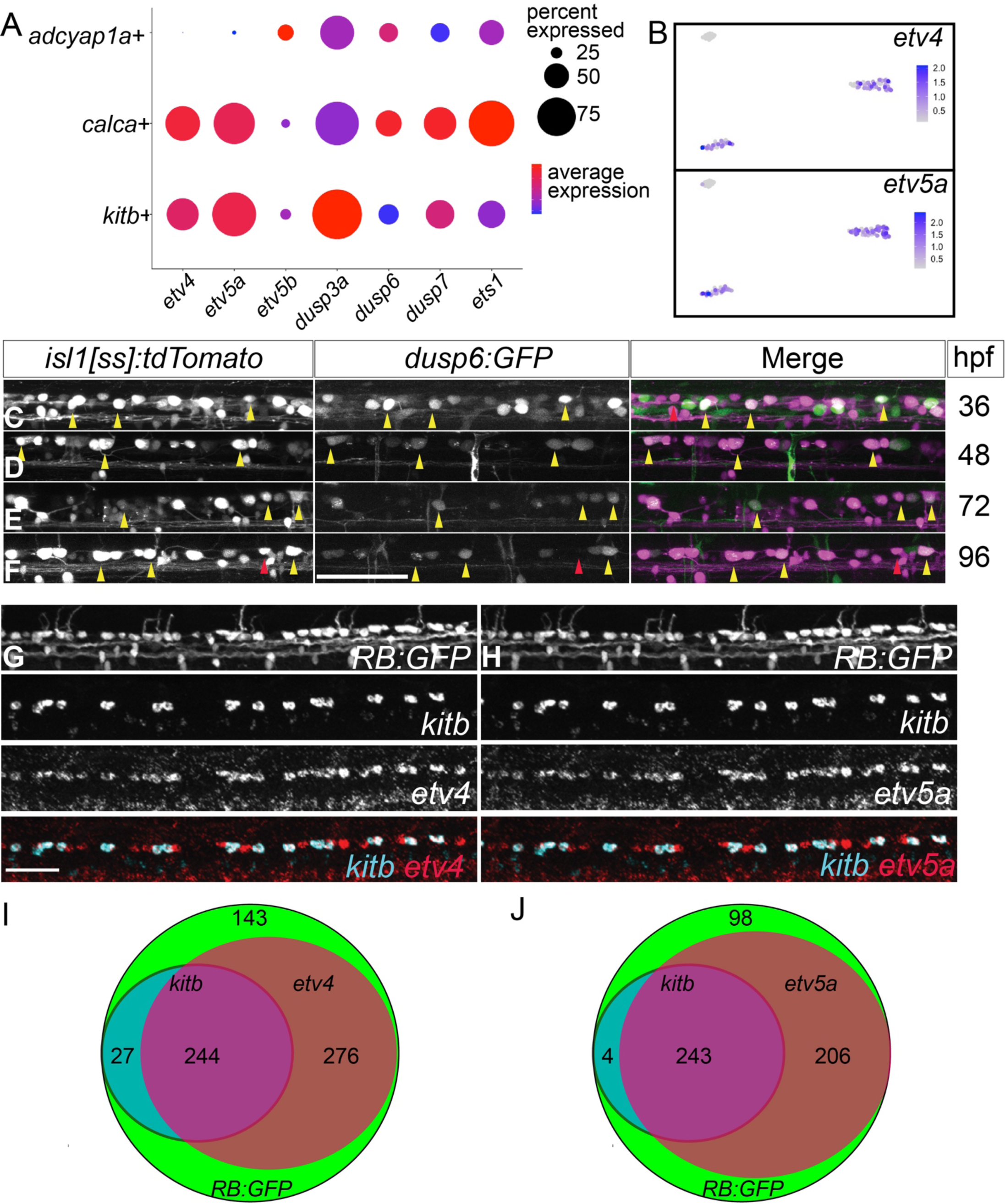
RBs express canonical downstream Fgf-signaling targets. (A) Dot plot shows expression of canonical Fgf targets in all three RB clusters. (B) Feature plots of two Fgf targets, *etv4* and *etv5a*, show expression in *kitb*+ and *calca*+ clusters. (C-F) Lateral views of dorsal neurons of live *isl1[ss]:tdTomato; dusp6:GFP* larvae at 36, 48, 72 and 96 hpf. Note that many RBs are GFP+ indicating active canonical Fgf signaling at these stages. (G,H) Lateral views of FISH staining in *RB:GFP* larvae for expression of *kitb* and downstream FGF signaling targets (G) *etv4* or (H) *etv5a*. Yellow arrowheads=*dusp6*+ RBs, Red arrowheads=*dusp6*-RBs, scale bars=50 μm. (I,J) Summed counts of RB markers in larvae with respective sets of FISH probes visualized in (G) and (H). The majority of RBs express *etv4* or *etv5a* and nearly all *kitb*+ RBs express *etv4*/*etv5a*.

### Fgf signaling is active in RB neurons

Our RB data set provided an opportunity to discover novel pathways that regulate development and maintenance of RB neurons. We are particularly interested in the role of receptor tyrosine kinase (RTK) signaling pathways, as these are major regulators of neural development, differentiation, and maintenance. In the case of RBs, the RTKs c-Kit and TrkC have critical roles in neuronal development and axon maintenance (Williams et al. 2000; Tuttle et al. 2022), but other important RTK pathways in this context have not been identified. Gene Ontology analysis (Ashburner et al. 2000) of all three RB subtypes for major RTK pathways revealed expression of genes in the FGF signaling pathway was increased 1.76 fold (number of observed divided by the number of expected genes in this GO category) in RBs. We found expression levels of *dusp*3a/6/7 and *ets1*, which are canonical Fgf targets, were increased in all three RB clusters (Fig. 4A). In contrast, other canonical Fgf targets were elevated in specific RB subclusters: *etv5b* in *adcyap1*+ cells, compared to *etv4* and *etv5a* in both *kit*+ and *calca*+ subclusters (Fig. 4A,B). This scRNA-seq expression data suggest that the Fgf pathway is active in most RB neurons.

First, to demonstrate active FGF signaling in RBs *in vivo*, we imaged transgenic larvae carrying a reporter of FGF signaling, *dusp6:eGFP* (Molina et al. 2007). This reporter transgene was introduced into a line carrying a driver/reporter system marking RBs, *isl1[ss]:LEXAV-P16;LEXAOP:tdTomato* (hereafter referred to as *isl1[ss]:RFP*)(Rasmussen et al. 2015). We observed EGFP signal from 36 hpf through 96 hpf in *isl1[ss]*+ RB neurons, indicating FGF signaling is active in most RBs (Fig. 4C-F). To verify the transcriptomic analysis, we assayed for RB-specific expression of two FGF downstream targets, *etv4* and *etv5a*, by FISH. These experiments showed that both transcripts are present in RB neurons (Fig. 4G,H). Quantitative analysis revealed they partially overlap with *kit*+ population as predicted by the scRNA-seq (Fig. 4I,J). Collectively, these data demonstrate active FGF signaling in RBs.

### Disrupting Fgf signaling in RB neurons leads to major loss of established axons

Fgf signaling has known important roles in neurogenesis and neuronal survival in the developing nervous system. But a role for Fgf signaling in functional, mature RBs has not been defined. To investigate this role, we disrupted Fgf signaling with pharmacological and genetic approaches in a temporally controlled manner. A seven-hour treatment with a pharmacological Fgf inhibitor, SU5402, substantially reduced expression of *dusp6:GFP* reporter in both *isl1[ss]*+ RB and motor neurons, indicating effective inhibition of Fgf signaling (Fig. 5A-B). Using a previously established protocol for characterizing the effect of RTK inhibitors on RB axons (Tuttle et al. 2022), we treated *RB:GFP* larvae beginning at 3 dpf with SU5402 for 48 h. This led to a major loss of innervation in the tail skin in a dose dependent manner (Fig. 5C-E). We then assessed the impact of temporal genetic loss of FGF signaling on RBs by combining the *isl1[ss]:RPF* transgene with a transgene for a heatshock-inducible dominant-negative FGFR (*hsp70:dnFGFR1a-GFP*) (Lee et al. 2005). This transgene has been extensively characterized in multiple studies to show inhibition of all four Fgf receptors (Nechiporuk et al. 2007; Wills et al. 2008; Konig and Jazwinska 2019; De Simone et al. 2021). Expression of *dnFgfr1a-GFP* following 30 min heat-shock at 72 hpf led to significant loss of RB axonal density in the tail at 4 dpf, a phenotype similar to SU5402 treatment (Fig. 5F-H). Combined, these data indicate a novel, critical role for Fgf signaling in maintaining established RB innervation of the skin.

**Figure 5:**
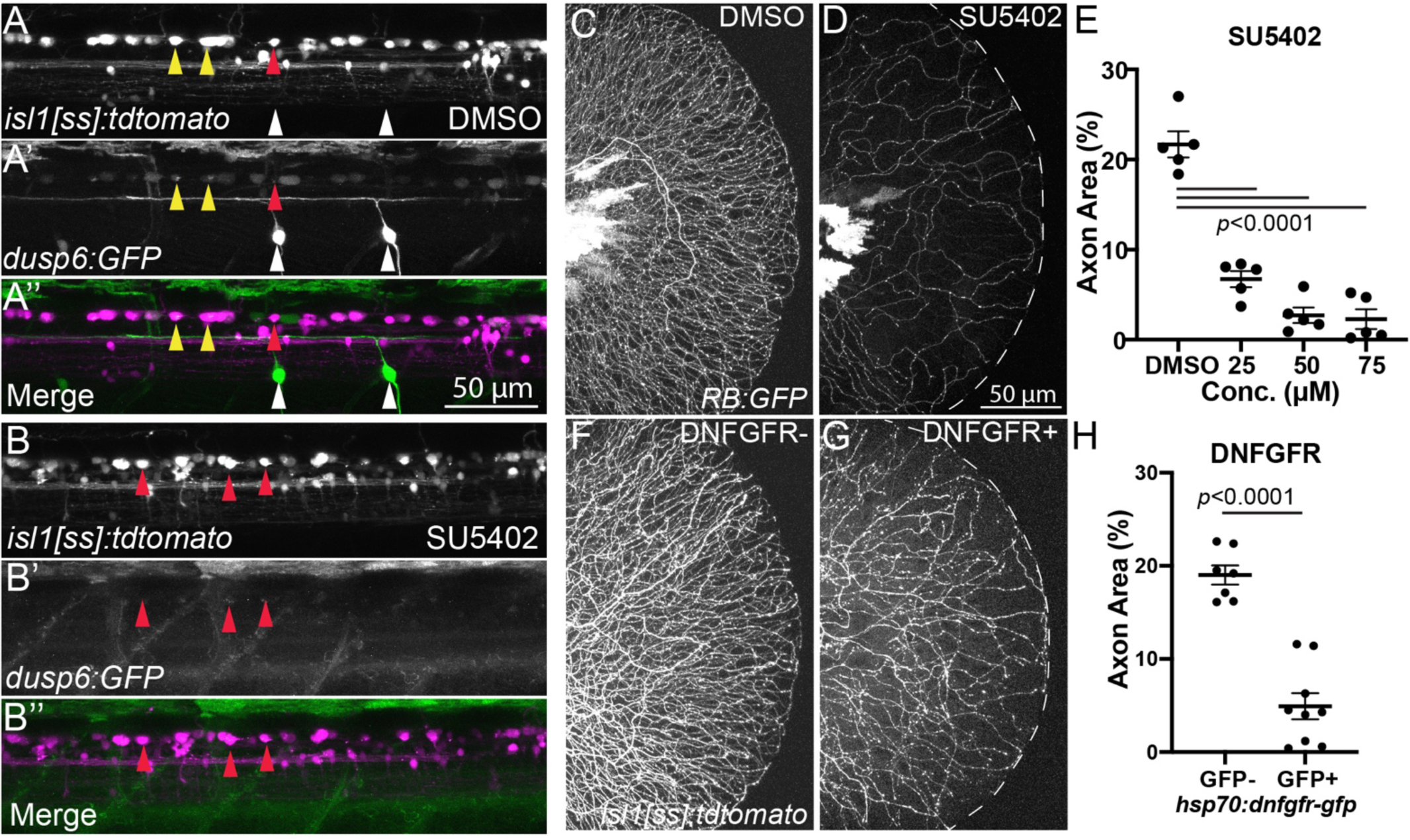
Inhibition of Fgf signaling induces loss of established sensory axons. (A,B) Lateral views of *isl1[ss]:tdTomato; dusp6:GFP* dorsal neurons in DMSO-(A) and SU5402-treated (B) 4 dpf larvae. Many RB neurons and motor neurons are GFP+ in DMSO-treated larvae but SU5402 treatment leads to loss of almost all GFP signal in dorsal neurons. Yellow arrowheads=GFP+ RB neurons, red arrowheads=GFP-RB neurons, white arrowheads=GFP+ motor neurons. (C,D) Lateral views of *RB:GFP* caudal tails at 5 dpf treated for 48 h with SU5402 (D), an Fgfr inhibitor, displayed substantial loss of RB axons in the tail compared to DMSO controls (C). (E) Quantification of the dose-dependent effect of SU5402 on tail axon density. DMSO=21.7±1.5%, 25 μM=6.7±0.9%, 50 μM=2.7±0.9%,75 μM=2.3±1.1%. (F,G) Lateral view of *isl1[ss]:tdTomato*+ control (F) and *isl1[ss]:tdTomato*; *hsp70:dn-FGFR1-GFP*+ caudal tails heatshocked at 4 dpf and imaged at 5 dpf. Compared to control (F), *hsp70:dn-FGFR1-GFP*+ larvae (G) had a reduction in RB axons similar to SU5402 treatment. (H) Quantification of tail axon-density following heat-shock. *Dnfgfr1-gfp*-=19.0±1.0%, *dnfgfr1-fgp*+=4.9±1.4%.

### FDA-approved drug that targets Fgfr induces somatosensory axon loss

Loss of somatosensory innervation of the skin underlies many peripheral neuropathies (Boyette-Davis et al. 2013; Fukuda et al. 2017), such as those caused by various cancer drugs. RTKs are a very common target of inhibitors used in precision cancer treatments in clinic, and inhibition of specific RTKs with targeted cancer drugs may underlie adverse somatosensory neurotoxicity (Roy et al. 2019; Tuttle et al. 2022). We asked if suppression of Fgf signaling with Fgfr inhibitors used in clinic may induce a similar neurotoxic phenotype to what we observe with pharmacological and genetic loss of Fgf signaling. Dovitinib is a potent Fgfr inhibitor that can also induce sensory peripheral neuropathies in patients (Loriot et al. 2010; Ma et al. 2019). Acute dovitinib treatment (7 hours) of *dusp6:GFP*;*isl1ss]:RFP* larvae led to loss of GFP signal in RBs and other neurons, a phenotype similar to the SU5402 treatment (Fig. 6A,B), indicating dovitinib significantly reduces RB Fgf signaling levels. To test if dovitinib-mediated inhibition of Fgfr impacted innervation of the skin, we treated *RB:GFP* larvae with dovitinib for 48 h and observed a dose-dependent, significant reduction of axonal density, consistent with the SU5402 phenotype (Fig. 6C-E). Taken together, these data show that suppression of Fgf signaling by a targeted cancer drug used in clinic may contribute to its sensory neurotoxicity.

**Figure 6:**
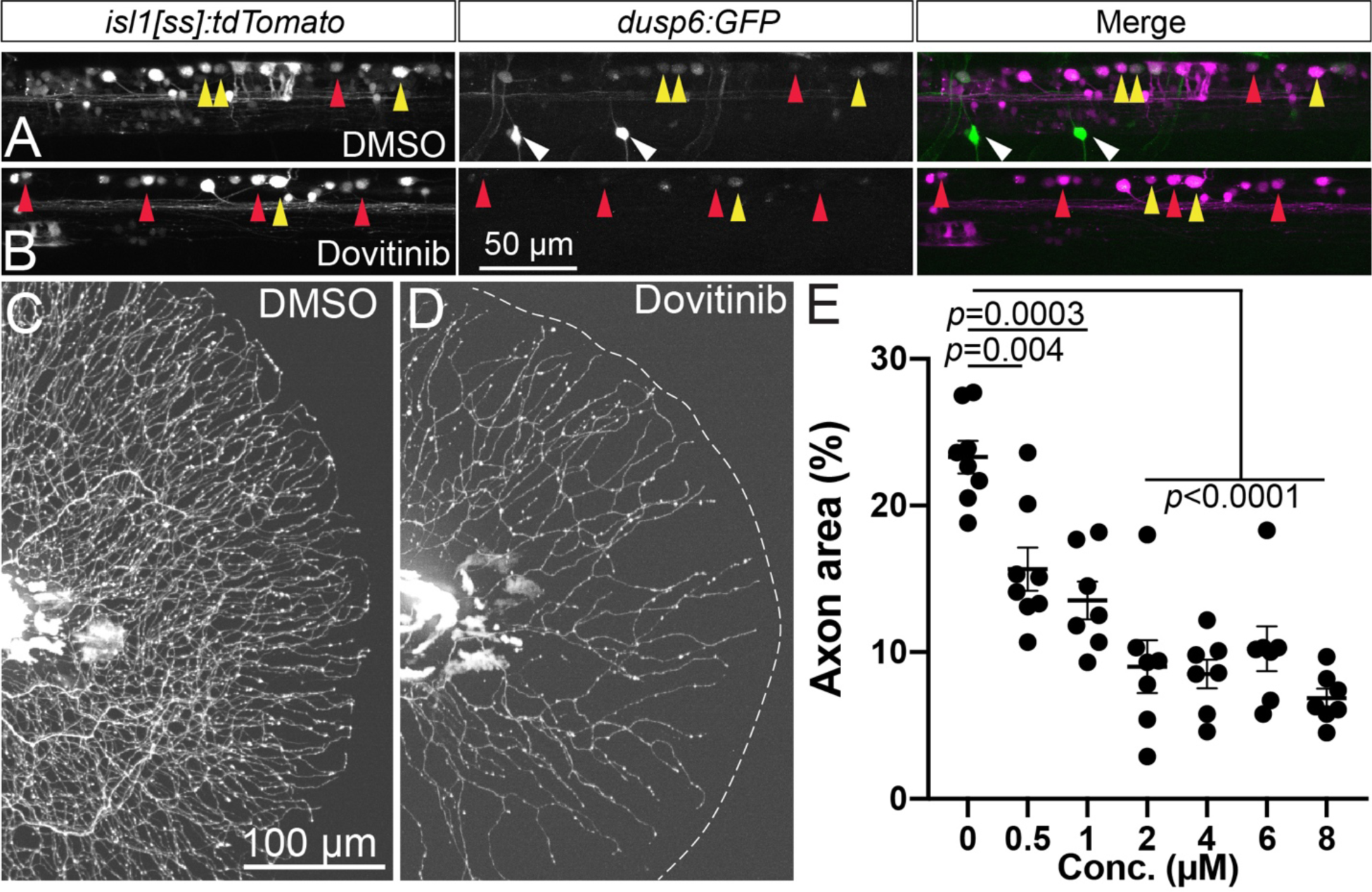
Dovitinib induces RB axon loss and reduction in Fgf signaling similar to SU5402 treatment. (A,B) Lateral view of *isl1[ss]:tdTomato; dups6:GFP* dorsal neurons in live 4 dpf larvae. Compared to DMSO-treated controls (A), short-term (7 h) dovitinib treatment (B) induced a substantial loss of Fgf signaling-dependent GFP expression. (C,D) Lateral view of *RB:GFP* larval caudal tails at 5 pf following 48 h DMSO (C) or dovitinib (D) treatment. Dovitinib treatment led to major axon loss similar to genetic and pharmacological loss of Fgfr signaling. (E) Quantification of dose-dependent effect of dovitinib on tail axon density. DMSO=23.3±1.1%, 0.5 μM=15.7±1.5%, 1 μM=13.5±1.3%, 2 μM=9.0±1.8%, 4 μM=8.5±1.0%, 6 μM=10.2±1.5%, 8 μM=6.9±0.6%. Yellow arrowheads=GFP+ RB neurons, red arrowheads=GFP-RB neurons, white arrowheads=GFP+ motor neurons.

### Wallerian-like degeneration and cell death underlies Fgf-mediated axon loss

We next explored the cellular bases of this dramatic loss of RB axons following inhibition of Fgf signaling. Two likely mechanisms could explain the axonal phenotype: axonal degeneration and RB cell death or axonal retraction. In order to distinguish between these possibilities, we performed timelapse imaging of *RB:GFP* neurons following treatment with SU5402 and dovitinib (Fig. 7 and Movies 1-6). After 24 h pretreatment with SU5402 (50 μM) a or dovitinib (6 μM), *RB:GFP*+ axons and cell bodies were imaged for ∼12 hours. In the vehicle controls, axons and their terminals remain for the most part stationary over time (yellow arrowheads in Fig. 7A and Movie 1). In contrast, both drug treatments caused Wallerian-like degeneration of RB axons over the course of 12 hours (magenta arrowheads in Fig. 7B,C and Movies 2,3). We did not observe any notable axon retraction for either SU5402 or dovitinib, although many axons remained stationary over the course of both treatments (Fig. 7B,C and Movies 2,3). Concurrent imaging during the same treatment period showed occasional disappearance of RB cell bodies, suggesting they undergo programmed cell death (Fig. 7D-F and Movies 4-6). To confirm whether an apoptotic cell death is the cellular basis of the phenotype, we immunolabeled SU5402- and dovitinib-treated larvae with antibodies that recognize cleaved Caspase-3. *RB:GFP* animals were incubated in the drug for 24 h then processed for the presence of cleaved Caspase-3 signal. We found a significant increase in Caspase3+ cells in SU5402- and dovitinib-treated animals (Fig. 7G-J). These data demonstrate that continued Fgf signaling is required in RB neurons for survival and/or axonal maintenance.

**Figure 7:**
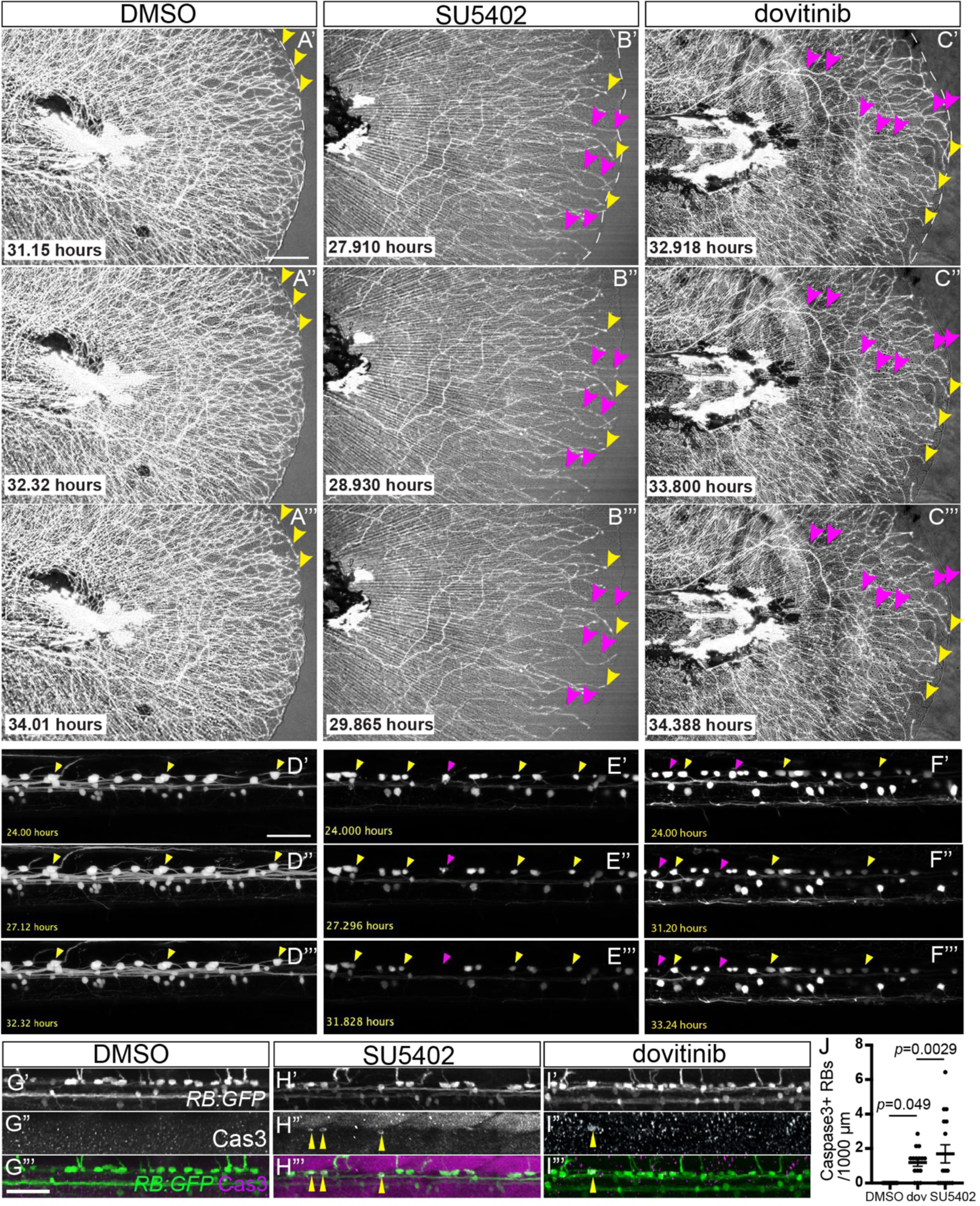
FGF inhibition and dovitinib treatment induce Wallerian-like axon degeneration and RB neuron apoptosis. (A-C) Stills from time-lapse movies of larval *RB:GFP* caudal tails treated with DMSO (A), SU5402 (B), or dovitinib (C). Yellow arrowheads=maintained axon terminals, magenta arrowheads=degenerating axons. (D-F) Stills from time lapse movies of larval *RB:GFP* dorsal neurons demonstrating RB apoptosis. Yellow arrowheads=surviving RB neurons, magenta arrowheads=RB neurons undergoing apoptosis. (G-I) Lateral view of *RB:GFP* dorsal neurons immunostained for cleaved Caspase-3 to mark apoptosis (yellow arrowheads). Compared to DMSO-treated larvae (G), SU5402 (H), and dovitinib (I). Scale bars=50 μm. (I) treatment displayed Cas3+ RB neurons. (J) Quantification of RB neuron apoptosis detected by Cas3+ immunostaining. DMSO=0±0 RB neurons, SU5402=1.2±.2, dovitinib=1.7±0.5. Scale bars=50 μm.

### Loss of Sarm1 activity rescues axonal degeneration caused by dovitinib

Next, we asked whether the degeneration of the cutaneous axons following dovitinib treatment can be mitigated by loss of Sarm1, a pro-degenerative protein (Osterloh et al. 2012). Sarm1 (Sterile Alpha and TIR Motif containing 1) is activated by nerve injury due to loss of nicotinamide adenine dinucleotide (NAD+) in damaged axons (Gerdts et al. 2015). Consequently, the loss of Sarm1 can block the degeneration of injured axons due to a number of drug insults (Geisler et al. 2016; Bosanac et al. 2021; Gould et al. 2021). To examine the role of Sarm1 following dovitinib treatment, we used a previously published loss-of-function, homozygous-viable allele of *sarm1* (Tian et al. 2020). We treated *RB:GFP*+ larvae derived from *sarm1* heterozygous incrosses with dovitinib or DMSO vehicle for 48 hours. Following the treatment, we examined axon density in *sarm1* homozygous mutants and their wild-type siblings. When treated with the DMSO, axon density of *sarm1* mutants were comparable to that of wild-type siblings (Fig. 8A,B,E). In contrast, axon density was significantly higher in *sarm1* mutants compare to sibs following dovitinib treatment (Fig 8C-E). These results show that the loss of Sarm1 suppresses the neurotoxic effect of dovitinib.

**Figure 8:**
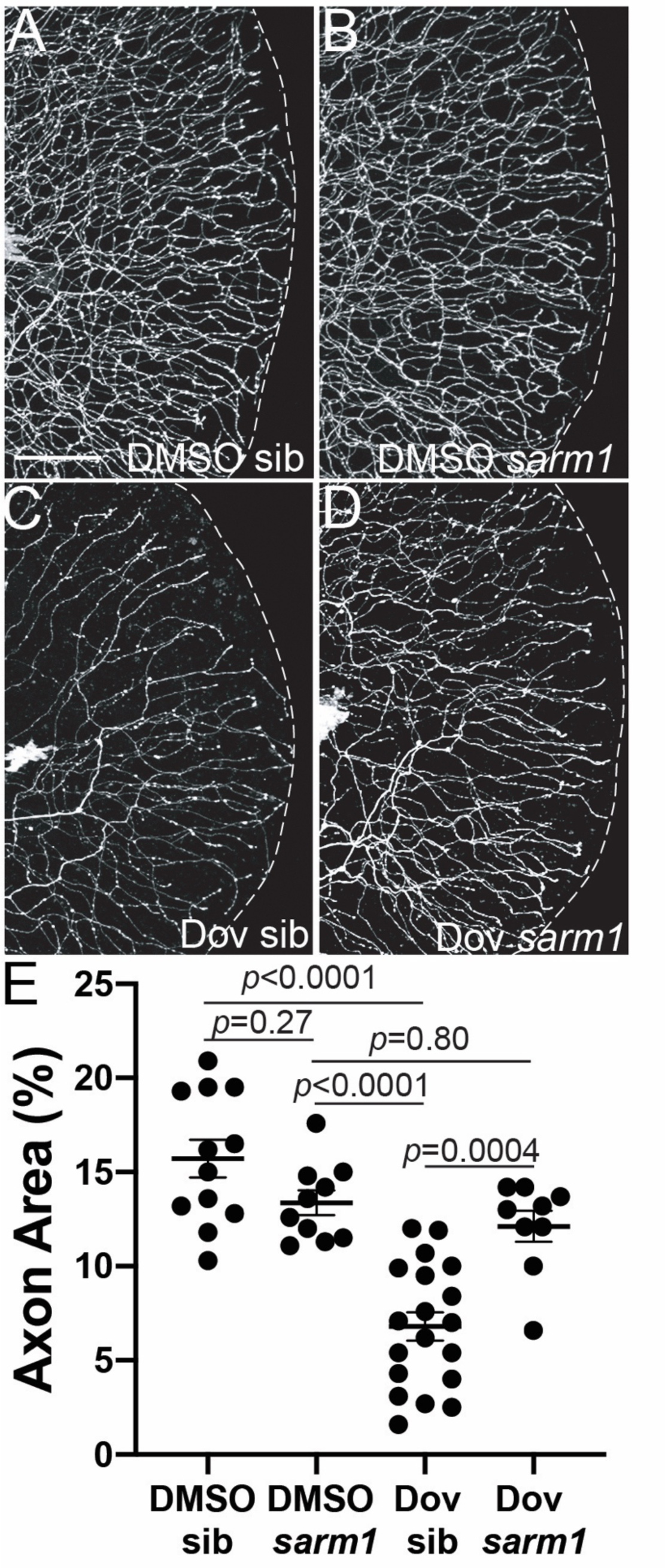
Sarm1 mediates dovitinib-induced axon degeneration. (A-D) Lateral views of *RB:GFP* caudal tails at 5 dpf of wild-type (A,C) or *sarm1-/-* mutants (B,D) treated with DMSO or dovitinib (3 - 5 dpf treatment). Compared to wildtype siblings treated with dovitinib, *sarm1* mutants had substantially higher GFP+ RB axon density. Scale bar=50 μm. (E) Quantification of axon density in dovitinib treatment of *sarm1* mutants. DMSO sib=15.7±1.0%, DMSO *sarm1* 13.4±0.7%, dovitinib sib=6.8±0.8%, dovitinib *sarm1*=12.1±0.8.

## DISCUSSION

Here, we provide a molecular characterization of RB neurons, a popular model for studies of sensory development and neurotoxicity. We show that on a transcriptional level, RB neurons can be separated into 3 distinct classes, which we designated *kit*+, *calca*+, and *adcyap1*+ populations. Based on single-molecule FISH, these are largely non-overlapping subpopulations present along the length of the spinal cord. Interestingly, only one RB population is clearly related to a specific subtype of TG or DRG sensory neurons in mice. Thus, it will require additional functional analysis to determine specific sensory modalities detected by each class of RB neurons. We also demonstrate that in zebrafish, RB neurons can be distinguished from TG neurons based on the expression of *hox* genes (head vs. trunk) as well as a small number of additional markers. Finally, we show the utility of our data set by investigating the requirement of the Fgf signaling pathway in RBs. Our data uncovers a new requirement for this pathway in axon maintenance and survival. Additionally, we find dovitinib, a small molecule drug that targets the Fgf pathway and is used in clinic to treat cancers, also induces RB degeneration. This RB axon degeneration can be prevented by the loss of Sarm1, suggesting an avenue for addressing neuropathic side effects of this drug. In summary, our data should provide a rich resource for investigating the molecular genetic mechanisms that guide functional diversification and maintenance of sensory neurons in vertebrates.

### Potential functions of RB subclasses

What are the potential functions of three distinct classes of RB neurons? We can speculate about this based on their similarity to known functional subclasses of mammalian TG and DRG neurons as well as presence of DE molecules (like channels and receptors) with known functions (underlined in Figs. 1K-M). As previously noted, *calca*+ RB neurons have high transcriptional similarity with group A fibers found in mammalian TG (NF3) and DRG (Aδ) neurons. These types of neurons function as cutaneous mechanoreceptors and nociceptors. In mammals, lightly myelinated Aδ-fiber peptidergic nociceptors express a variety of neuropeptides including *Calca* and *Tac1*. However, *tac1* is only expressed at low levels in *calca*+ RBs. Nevertheless, consistent with this functional homology, *ntrk2* is highly expressed in both *calca*+ RBs and mammalian Aδ neurons. In summary, the transcriptional profile of *calca*+ RB neurons is consistent with them functioning as mechanoreceptors and nociceptors. However, only partial similarity to NF3- and Aδ-subtypes suggests that *calca*+ RBs mediate additional sensory modalities.

Interestingly, while *kitb*+ neurons display a weak homology to NP and PEP TG and DRG neurons, their top differentially upregulated genes like *prkcaa*, *p2rx3a/b*, and *trpc4a*, argue that these neurons detect distinct nociceptive somatosensory stimuli. Notably, Trpc4 mediates itch sensation in mammals (Bandell et al. 2004). Htr3a, a serotonergic receptor involved in response to itch (Ostadhadi et al. 2015), is also highly expressed in *kitb*+ neurons. Adult zebrafish demonstrate pruritic behaviors (Esancy et al. 2018); thus it is possible that this class of RB neurons mediates itch or a similar modality in larval and juvenile fish as well. Another Trp channel, which is highly expressed in *kitb*+ RB neurons, is *trpa1b*. In mammalian DRGs, *Trpa1* is expressed in a subset of PEP neurons and is a multiple irritant sensor, including mustard oil (Story et al. 2003; Bandell et al. 2004; Jordt et al. 2004). Loss-of-function study in zebrafish supports the role of Trpa1 channel in chemical (e.g. mustard oil) but not thermal or mechanical sensation (Prober et al. 2008). In addition, the Trpa1 channel has been implicated in pruritic like behavior in mammals (Wilson et al. 2011) as well as adult zebrafish (Esancy et al. 2018). In summary, *kitb*+ neurons are likely mutlimodal, sensing both mechanical stimuli akin to itch via Trpa1, Trpc4 and Htr3a channels as well chemical irritants via the Trpa1 channel.

Finally, RB neurons of *adcyap1a*+ subtype are not transcriptionally similar to any specific subtypes of mammalian TG or DRG sensory neuron. In mammalian DRG, high levels of Adcyap1 mark PEP1.1 and PEP1.2 subtypes (Jung et al. 2023). These are homologous to PEP TG neurons, but neither displays any transcriptional similarity to *adcyap1a*+ RB neurons. Thus, additional functional analysis is necessary to assign them a specific sensory role.

### Role of Fgf signaling in RB survival and its potential role in DIPNs

Fgf signaling plays a number of essential roles during nervous system development, including morphogenesis and patterning, neuroblast proliferation and survival, and axonal pathfinding. However, the role for this pathway in nervous system maintenance is less well understood, although Fgf activation does have a neuroprotective role following CNS injury (Klimaschewski and Claus 2021). Here, we demonstrate a new role for the Fgf signaling pathway in mature RB axon maintenance and cell survival. In the absence of Fgf signaling, RB axons undergo Wallerian degeneration concurrently with programmed cell death. Importantly, treatment with a multi-kinase inhibitor (MKI), dovitinib, an FDA-approved Fgf receptor inhibitor, produced a phenotype very similar to both pharmacological and genetic inhibition of the Fgf pathway. Presence of multiple canonical Fgf targets in RB neurons as well as expression of Fgf reporter argue that Fgf signaling is required for RB maintenance. We noted that two Fgf factors, Fgf12b and Fgf13a/b, are expressed in all RB neurons. Fgf11-14 are referred to as Fgf homologous factors, because despite having a structural similarity with secreted Fgf ligands they are generally believed to function as intracellular, non-secreted molecules. In neurons, these factors can act as modulators of voltage-gated channels (Yan et al. 2013; Singh et al. 2021). In fact, interaction of Fgf13 with the sodium channel Na_v_1.7 is required for noxious heat sensation by DRG neurons (Yang et al. 2017). However, a recent study demonstrates that Fgf homologous factors can be secreted into extracellular space (Biadun et al. 2023). In this case, they signal through the canonical Fgf signaling pathway involving biding of Fgfr1 and activation of Erk1/2. On a cellular level, activation of this particular Fgf pathway is necessary to protect cells from apoptosis. Given high expression of Fgf12b and Fgf13a/b in RB neurons, it is tempting to speculate that this autocrine mechanism is necessary for RB cell survival. Although, we cannot exclude that canonical Fgfs ligands are secreted by the surrounding tissues to promote Fgf activation in RB maintenance.

Finally, we demonstrate that dovitinib, an MKI that is currently used in the clinic to treat multiple cancers, causes the Wallerian degeneration of RB axons and RB cell death. MKIs, like dovitinib, are a relatively new and effective class of anti-cancer therapies. However, many MKIs, including dovitinib, cause unintended neurotoxic effects. We found that the dovitinib-induced axonal loss can be suppressed by the absence of Sarm1. This finding may provide a therapeutic opportunity to address these painful side effects in the clinic and can potentially be extended to other MKIs that have similar neurotoxic effects.

## MOVIES

Movie 1: Time-lapse imaging of RB axons during control DMSO treatment between ∼96 and 108 hpf. Each time frame is a Z projection from the whole tail.

Movie 2: Time-lapse imaging of RB axons during SU5402 treatment between ∼96 and 108 hpf. Each time frame is a Z projection from the whole tail.

Movie 3: Time-lapse imaging of RB axons during dovitinib treatment between ∼96 and 108 hpf. Each time frame is a Z projection from the whole tail.

Movie 4: Time-lapse imaging of RB cell bodies during control DMSO treatment between ∼96 and 108 hpf. Each time frame is a Z projection RB cell bodies from one side of the animal.

Movie 5: Time-lapse imaging of RB cell bodies during SU5402 treatment between ∼96 and 108 hpf. Each time frame is a Z projection RB cell bodies from one side of the animal.

Movie 6: Time-lapse imaging of RB cell bodies during dovitinib treatment between ∼96 and 108 hpf. Each time frame is a Z projection RB cell bodies from one side of the animal.

## AUTHOR CONTRIBUTIONS

Conceived and designed experiments: AT and AN. Performed scRNA-seq experiments: LM, HW, and JK. Performed remaining experiments: AT, LR, and AN. Analyzed data for scRNA-seq: NC, LH, LM, and AN. Analyzed remaining data: AT, LR, LM, and AN. Wrote Manuscript: AT and AN.

## ACKNOWLEDGEMENTS

The authors thank Drs. Hernán Lopez-Schier, Jeff Rasmussen, and Kara Cerveny for reagents. We also thank Dr. Kelly Monk for generous access to her VAST sorting and confocal imaging system, and Dr. Kevin Wright for comments on the manuscript. This work was supported with funding provided to AVN from the NCI (R21 CA260025; http://www.nci.nih.gov) and the NINDS (R01 NS111419; http://www.ninds.nih.gov) as well as to Dr. Paul Brehm’s laboratory at the Vollum Institute (HW and JK) from the NINDS (NS105664).

